# Video-based quantification of human movement frequency using pose estimation

**DOI:** 10.1101/2021.02.01.429161

**Authors:** Hannah L. Cornman, Jan Stenum, Ryan T. Roemmich

## Abstract

Assessment of repetitive movements (e.g., finger tapping) is a hallmark of motor examinations in several neurologic populations. These assessments are traditionally performed by a human rater via visual inspection; however, advances in computer vision offer potential for remote, quantitative assessment using simple video recordings. Here, we evaluated a pose estimation approach for measurement of human movement frequency from smartphone videos. Ten healthy young participants provided videos of themselves performing five repetitive movement tasks (finger tapping, hand open/close, hand pronation/supination, toe tapping, leg agility) at four target frequencies (1-4 Hz). We assessed the ability of a workflow that incorporated OpenPose (a freely available whole-body pose estimation algorithm) to estimate movement frequencies by comparing against manual frame-by-frame (i.e., ground-truth) measurements for all tasks and target frequencies using repeated measures ANOVA, Pearson’s correlations, and intraclass correlations. Our workflow produced largely accurate estimates of movement frequencies; only the hand open/close task showed a significant difference in the frequencies estimated by pose estimation and manual measurement (while statistically significant, these differences were small in magnitude). All other tasks and frequencies showed no significant differences between pose estimation and manual measurement. Pose estimation-based detections of individual events (e.g., finger taps, hand closures) showed strong correlations with manual detections for all tasks and frequencies. In summary, our pose estimation-based workflow accurately tracked repetitive movements in healthy adults across a range of tasks and movement frequencies. Future work will test this approach as a fast, low-cost, accessible approach to quantitative assessment of repetitive movements in clinical populations.

## INTRODUCTION

Several neurologic conditions – including Parkinson’s disease (PD; 1) and cerebellar disorders (2) – impair the ability to execute repetitive movements. This has led to the inclusion of repetitive movements in many clinical examinations of motor function. For example, the MDS-UPDRS (the clinical standard for motor assessment in PD) contains several items that test repetitive movements across different tasks, including finger tapping, hand opening and closing, hand pronation and supination, toe tapping, and leg agility (3). These assessments provide important insights into disease severity that influence treatment decisions and enable tracking of disease progression over time.

Traditional assessments of repetitive movements rely on subjective clinical scales rated by a trained clinician. For example, when performing Part III of the MDS-UPDRS (3), a neurologist rates the performance of each repetitive movement task from 0 (no impairment) to 4 (severe impairment) via visual assessment. Other approaches have used research-grade equipment (e.g., motion capture systems), wearables (4,5), and mobile devices (6–8) to quantify performance of repetitive movements. However, motion capture systems are inaccessible to many clinicians and researchers, and wearable- or smartphone-based approaches typically assess only specific movements (e.g., finger tapping). There is a need for a rapid, low-cost approach capable of providing quantitative measurement of repetitive movements across multiple extremities using equipment that is readily accessible in the home or clinic.

Recent developments in computer vision have produced pose estimation algorithms capable of tracking human movement with a high degree of accuracy and minimal technological requirement (9–14). These algorithms have shown an impressive ability to track specific features of the human body (e.g., fingers, leg joints) from digital videos recorded by common household devices (e.g., smartphones, tablets, laptop computers). There is significant potential for these approaches to provide rapid, automated measurement and analysis of repetitive movements from videos recorded in the home or clinic, enabling remote assessment that does not require a patient visit to the clinic or manual assessment from a clinician.

In this study, we used OpenPose (a freely available human pose estimation algorithm (11,15,16) to measure the frequencies of repetitive upper and lower extremity movements using smartphone videos. Previous studies have used OpenPose to study human gait (17–21) and quantify tremor severity in persons with PD (22). Other pose estimation algorithms have been used to measure the degree of levodopa-induced dyskinesias (23) and finger tapping frequency in PD that correlated well with clinical ratings (24,25). While emerging evidence suggests significant potential for tracking repetitive movements using video-based pose estimation approaches, here we addressed two important unmet needs: 1) testing across a range of clinically relevant tasks spanning the upper and lower extremities, and 2) assessment of accuracy via validation against ground-truth measurement. Based on our prior work (21), we expected good agreement between the OpenPose estimates of movement frequency and ground-truth measurement. We also share our perspectives regarding limitations, potential problems, and suggestions for successful analysis. All code used for this project is freely available at https://github.com/RyanRoemmich/PoseEstimationApplications.

## MATERIALS AND METHODS

### Participants

We recruited ten healthy adults (four male, six female, age (mean ± standard deviation): 28 ± 4 years) to participate. All participants provided written informed consent in accordance with the Johns Hopkins University School of Medicine Institutional Review Board.

### Data collection

Participants provided digital videos of themselves performing five repetitive movement tasks included on the MDS-UPDRS (3): finger tapping, hand opening and closing, hand pronation and supination, toe tapping, and leg agility (i.e., heel tapping). We asked participants to perform each task at four target frequencies (1 Hz, 2 Hz, 3 Hz, and 4 Hz) using an online metronome for approximately 10 seconds each. Each participant received instructions for filming the videos and we provided example videos of each task to promote uniformity in the recordings. The instructions were as follows:

#### Setup

- Sit in a chair with knees pointing forward and feet flat on the floor (with the exception of hand pronation/supination, where we asked participants to record sagittal views – “record from the side” – of themselves performing the movement).
- Make sure hand is held to the side when performing hand and finger tasks, not in front of the body.
- The phone should be approximately five feet from the participant and at their eye level when filming.
- Capture the whole body.
- Film with a smartphone camera at standard sampling rate (approximately 30 frames/sec).
- Use an online metronome to pace movement frequency and try to match the metronome as closely as possible.

#### Video recording

- Record approximately 10 seconds of the participant performing each of the following tasks with one hand at 4 different speeds (1-4 Hz).
- *<u>Finger tapping</u>*: tap index finger to the thumb (opening as big as possible between taps).
- *<u>Hand opening and closing</u>*: fully open and close the hand repeatedly (opening as big as possible between hand closures).
- *<u>Hand pronation and supination</u>*: alternate pronating and supinating the hand (pronate on one beat, supinate on the next).
- *<u>Toe tapping</u>*: keep the heel planted and tap the toe to the ground.
- *<u>Leg agility</u>*: lift the entire foot and strike the ground with heel.

### Pose estimation

Our data analysis workflow is shown in Figure 1. First, we analyzed video recordings in OpenPose using a Python environment to obtain two-dimensional pixel coordinates of OpenPose keypoints. We used both the Hand (26) and BODY_25 (16) keypoint models simultaneously. The Hand keypoint model tracks 21 keypoints on each hand: one on each tip and all three joints per finger, and one at the base of the hand. The BODY_25 keypoint model tracks keypoints at the nose, neck, midhip, and bilateral eyes, ears, shoulders, elbows, wrists, hips, knees, ankles, heels, big toes, and small toes. The outputs of the OpenPose analyses yielded: 1) JSON files containing frame-by-frame pixel coordinates of each keypoint, and 2) an output video file in which a skeleton figure representing the detected keypoints is overlaid onto the original video recording. We then analyzed the JSON files using custom MATLAB (Mathworks, Natick, MA) software to calculate movement frequency (detailed below).

**Figure 1.**
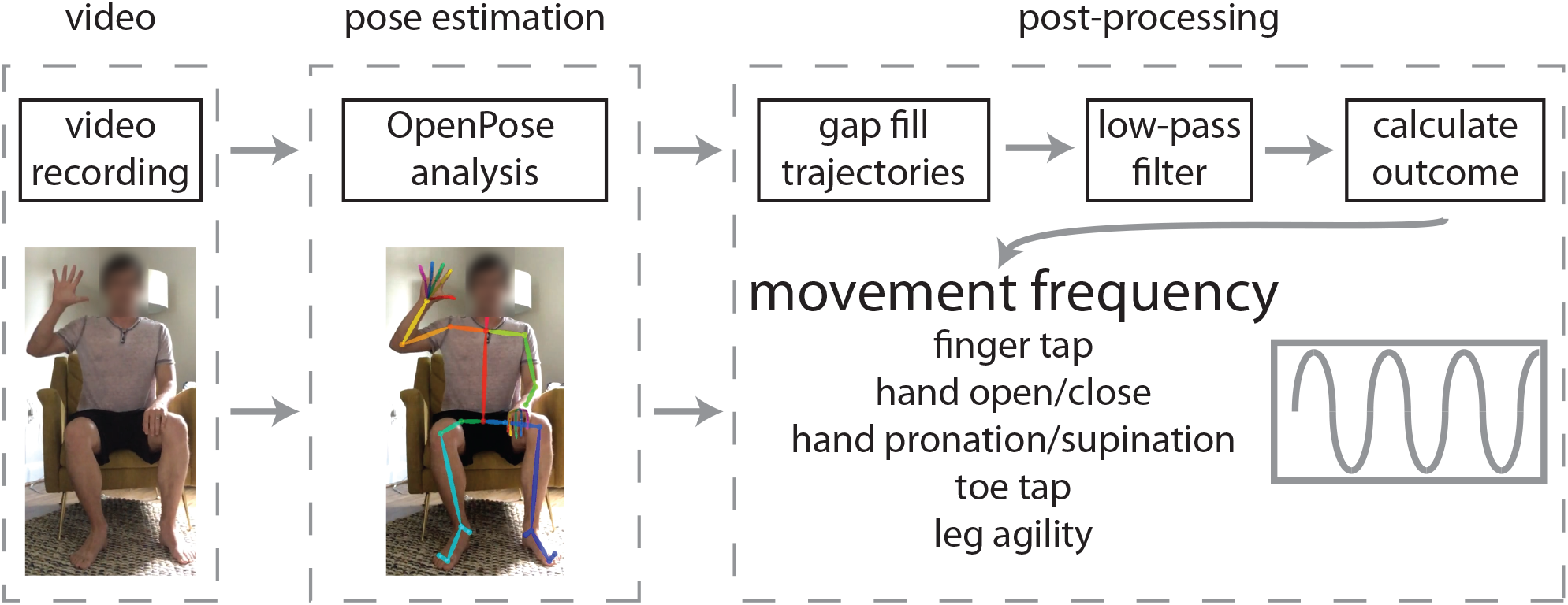
We used a three-step workflow to measure movement frequency from smartphone videos. Participants first provided a video recording of themselves performing the task of interest (we assessed five different tasks in this study: finger tapping, hand opening/closing, hand pronation/supination, toe tapping, and leg agility or heel tapping). Next, we used OpenPose to estimate body and hand keypoint locations on each video. We then developed post-processing software to calculate movement frequency using the OpenPose outputs.

### Calculation of movement frequency

We performed several steps to calculate movement frequencies from the JSON files. First, we filled gaps in all keypoint trajectories (i.e., frames where OpenPose did not detect all keypoints) using linear interpolation. Second, we filtered keypoint trajectories using a zero-lag 4^th^ order low-pass Butterworth filter with a cut-off frequency at 5 Hz. Third, we identified events (finger taps, hand closures, hand pronations/supinations, toe taps, heel taps) using the following criteria:

a. *Finger tapping* and *hand opening and closing* tasks: local minima in the distance between index finger and thumb keypoints.
b. *Hand pronation and supination*: local maxima (thumb up, supination) and minima (thumb down, pronation).
c. *Toe tapping*: local minima of the big toe keypoint.
d. *Leg agility*: local minima of the heel keypoint.

Finally, we calculated movement frequency from the identified event times.

### Manual ground-truth identification of event times

We obtained a ground-truth benchmark by manually identifying event times through frame-by-frame visual inspection for all videos. We used the same criteria listed above to identify events manually. From these manually-identified event times, we calculated ground-truth movement frequency.

### Statistical analysis

We compared movement frequencies between methods and among target frequencies using a 2×4 method (pose estimation, manual) x target frequency (1, 2, 3, 4 Hz) repeated measures ANOVA. All events identified by the pose estimation and manual methods were included in calculations of movement frequencies.

For all correlational analyses, we isolated events that had been detected by both the pose estimation and manual identification methods (the overwhelming majority of events were detected by both methods, as described in the Results section below). To do this, we first compared the number of events identified by both methods in each individual video. If the numbers of events were not equal between the two methods (i.e., either the pose estimation approach identified more events than the manual identification or vice versa), we discarded the events from the method with more detected events that had video timestamps (“event times”) most dissimilar to any event time detected by the other method until the number of detected events were equivalent in both methods. Most often, this meant that the pose estimation approach detected (or failed to detect) an additional event at the beginning or end of the trial, and these events that were not detected by both methods were then discarded from the correlational analyses. We also observed a few trials where both methods detected the same number of events, but one method had detected an additional event at the beginning of the trial and the other an additional event at the end of the trial; in this case, we discarded both unmatched events. From the matched events, we calculated Pearson correlation coefficients (*r*) and intra-class correlation coefficients (ICC_C-1_ and ICC_A-1_) to assess correlations (*r*), consistency (ICC_C-1_) and agreement (ICC_A-1_) between event time estimates by our pose estimation approach and those identified manually. We used an alpha level of 0.05 for all analyses and performed post-hoc pairwise comparisons with Bonferroni corrections where appropriate.

## RESULTS

### Finger tapping

Repeated measures ANOVA revealed a significant main effect of target frequency (F(3,27)=416.31, p<0.001) but no significant main effect of method (F(1,9)=1.43, p=0.26) or target frequency x method interaction (F(3,27)=0.97, p=0.42) on frequency of finger tapping (Figure 2A). As expected, post-hoc analyses revealed significant differences in movement frequency between each pair of target frequencies (all p<0.001). This confirmed that participants changed their tapping frequency in accordance with the metronome-paced target frequencies, and these changes were similar in the pose estimation and ground-truth measurements.

**Figure 2.**
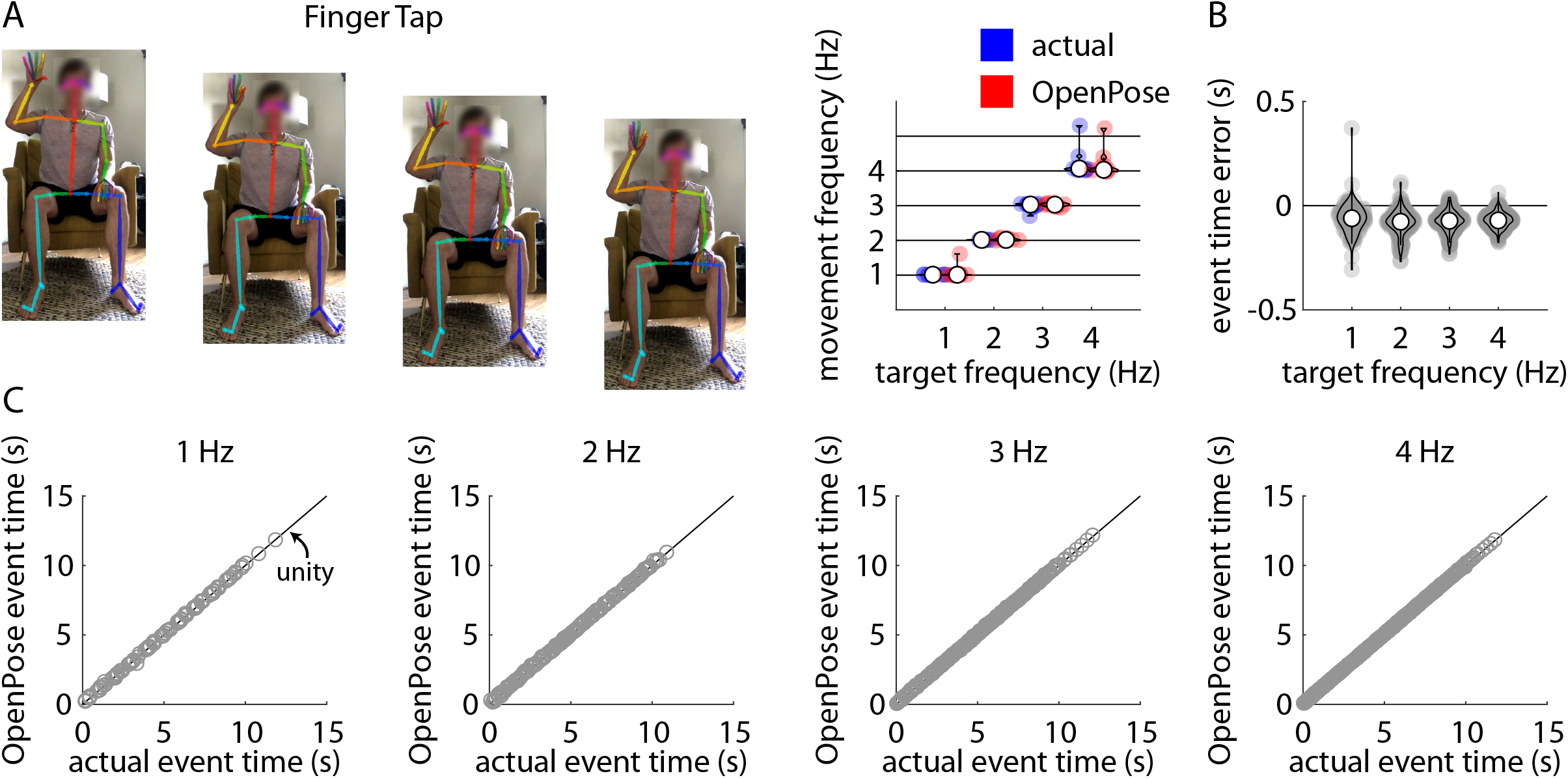
A) Representative example of stillframes from an OpenPose output video of a participant performing the finger tapping task (left) and violin plots showing comparisons of individual participant mean finger tapping frequencies measured manually (“actual”, blue) and using our pose estimation workflow (“OpenPose”, red) across all four target frequencies (right). B) Violin plots showing event time errors (i.e., differences between video timestamps of finger taps as measured manually and using pose estimation) for each individual finger tap across all participants. In Figures 2A and 2B, white circles indicate group mean values, shaded circles indicate individual data points. These conventions are consistent across Figures 2–6. C) Scatter plots showing relationships between the video timestamps of all individual finger taps across all participants as measured manually (x-axis) or using our pose estimation workflow (y-axis) across all four target frequencies. Relevant correlation coefficients are included in the Results section of the text for Figures 2–6.

Across all trials, our pose estimation approach identified 993 finger taps and manual identification detected 1002 finger taps across all participants and trials; of these, there were 986 common finger taps detected by both methods. Distributions of errors between the pose estimation and manual detections of each finger tap (i.e., time of manually-detected finger tap minus time of pose estimation-detected finger tap) for all taps detected by both methods are shown in Figure 2B. Figure 2C shows associations between manually-detected finger tap times and pose estimation-detected finger tap times for target frequencies of 1 Hz (*r*=1.00, p<0.001; ICC_C-1_=1.00, p<0.001; ICC_A-1_=1.00, p<0.001), 2 Hz (*r*=1.00, p<0.001; ICC_C-1_=1.00, p<0.001; ICC_A-1_=1.00, p<0.001), 3 Hz (*r*=1.00, p<0.001; ICC_C-1_=1.00, p<0.001; ICC_A-1_=1.00, p<0.001), and 4 Hz (*r*=1.00, p<0.001; ICC_C-1_=1.00, p<0.001; ICC_A-1_=1.00, p<0.001).

### Hand opening and closing

Repeated measures ANOVA revealed a significant main effect of target frequency (F(3,27)=1032.07, p<0.001), a significant main effect of method (F(1,9)=9.08, p=0.02), and a significant target frequency x method interaction (F(3,27)=4.70, p=0.01) on frequency of hand opening and closing (Figure 3A). Post-hoc analyses revealed significant differences in movement frequency between each pair of target frequencies (all p<0.001). The movement frequencies estimated by the pose estimation approach were significantly slower than the frequencies identified manually at target frequencies of 2 Hz, 3 Hz, and 4 Hz (all p<0.02); however, the magnitudes of these differences were very small (mean difference between methods no larger than 0.03 Hz for any target frequency).

**Figure 3.**
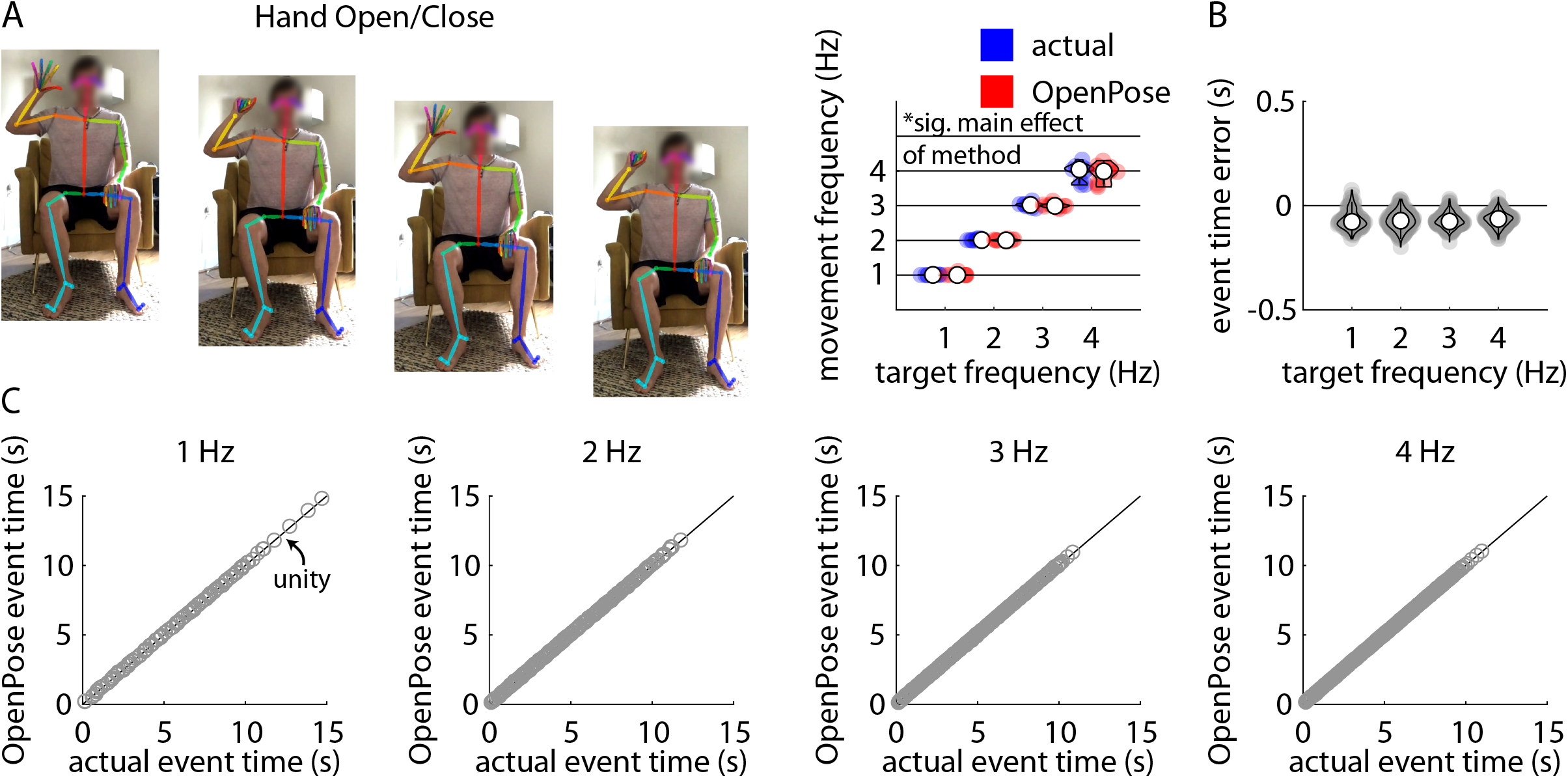
A) Representative example of stillframes from an OpenPose output video of a participant performing the hand opening and closing task (left) and violin plots showing comparisons of individual participant mean hand closure frequencies measured manually (“actual”, blue) and using our pose estimation workflow (“OpenPose”, red) across all four target frequencies (right). B) Violin plots showing event time errors (i.e., differences between video timestamps of hand closures as measured manually and using pose estimation) for each individual hand closure across all participants. C) Scatter plots showing relationships between the video timestamps of all individual hand closures across all participants as measured manually (x-axis) or using our pose estimation workflow (y-axis) across all four target frequencies.

Across all trials, our pose estimation approach identified 960 hand closures and manual identification detected 973 hand closures across all participants and trials; of these, there were 955 hand closures detected by both methods. Distributions of errors between the pose estimation and manual detections of each hand closure detected by both methods are shown in Figure 3B. Figure 3C shows associations between manually-detected hand closure times and pose estimation-detected hand closure times for target frequencies of 1 Hz (*r*=1.00, p<0.001; ICC_C-1_=1.00, p<0.001; ICC_A-1_=1.00, p<0.001), 2 Hz (*r*=1.00, p<0.001; ICC_C-1_=1.00, p<0.001; ICC_A-1_=1.00, p<0.001), 3 Hz (*r*=1.00, p<0.001; ICC_C-1_=1.00, p<0.001; ICC_A-1_=1.00, p<0.001), and 4 Hz (*r*=1.00, p<0.001; ICC_C-1_=1.00, p<0.001; ICC_A-1_=1.00, p<0.001).

### Hand pronation and supination

Repeated measures ANOVA revealed a significant main effect of target frequency (F(3,27)=656.54, p<0.001), but no significant main effect of method (F(1,9)=0.33, p=0.58) or target frequency x method interaction (F(3,27)=2.15, p=0.12) on frequency of alternating hand pronation and supination (Figure 4A). Post-hoc analyses revealed significant differences in movement frequency between each pair of target frequencies (all p<0.001).

**Figure 4.**
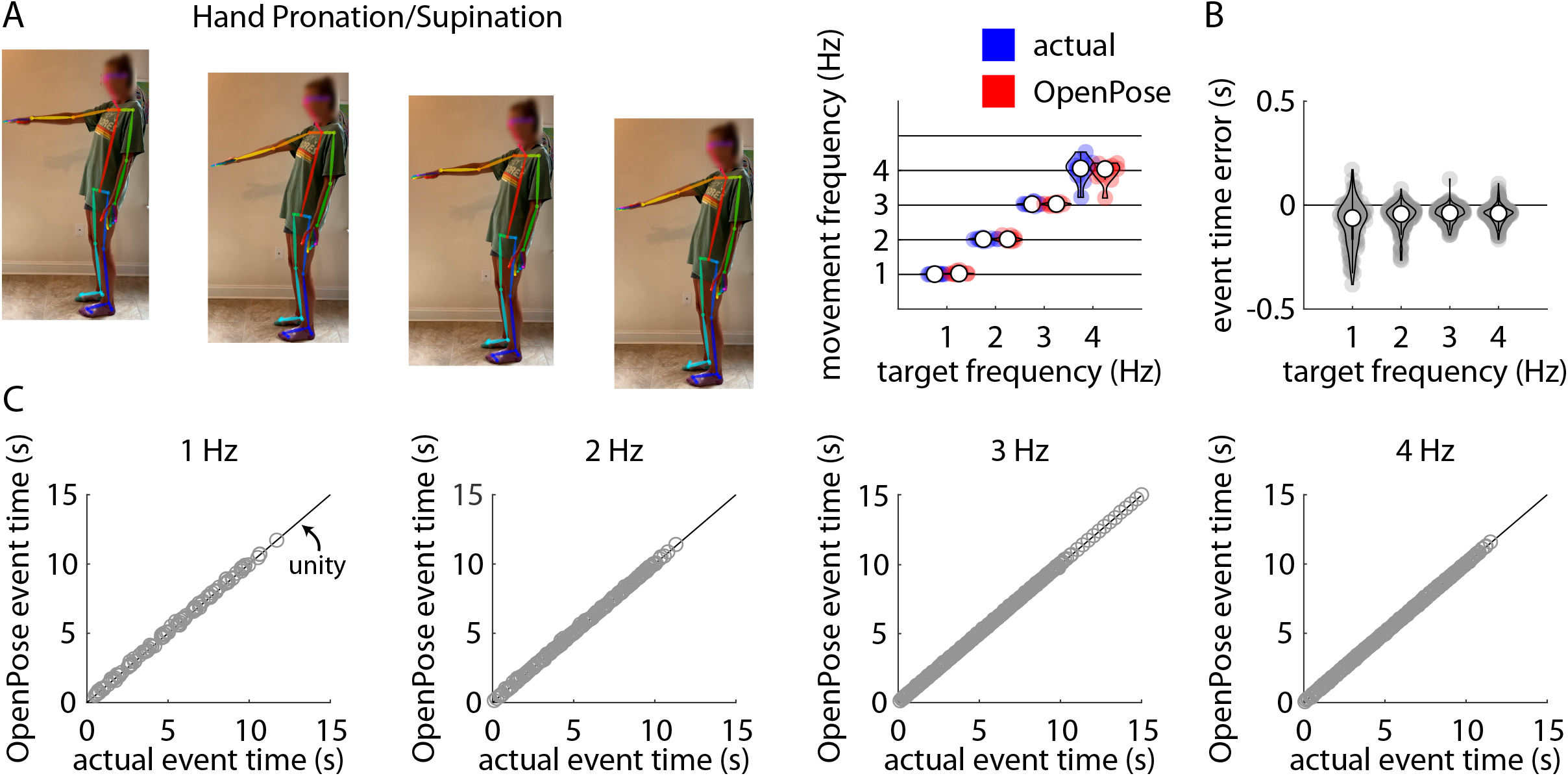
A) Representative example of stillframes from an OpenPose output video of a participant performing the hand pronation/supination task (left) and violin plots showing comparisons of individual participant mean hand pronation/supination frequencies measured manually (“actual”, blue) and using our pose estimation workflow (“OpenPose”, red) across all four target frequencies (right). B) Violin plots showing event time errors (i.e., differences between video timestamps of hand pronation or supination events as measured manually and using pose estimation) for each individual hand pronation or supination event across all participants. C) Scatter plots showing relationships between the video timestamps of all individual hand pronation or supination events across all participants as measured manually (x-axis) or using our pose estimation workflow (y-axis) across all four target frequencies.

Across all trials, our pose estimation approach identified 974 hand pronation or supination events and manual identification detected 991 hand pronation or supination events across all participants and trials; of these, there were 958 common hand pronation or supination events detected by both methods. Distributions of errors between the pose estimation and manual detections of each hand pronation or supination event detected by both methods are shown in Figure 4B. Figure 4C shows associations between manually-detected hand pronation or supination event times and pose estimation-detected hand pronation or supination event times for target frequencies of 1 Hz (*r*=1.00, p<0.001; ICC_C-1_=1.00, p<0.001; ICC_A-1_=1.00, p<0.001), 2 Hz (*r*=1.00, p<0.001; ICC_C-1_=1.00, p<0.001; ICC_A-1_=1.00, p<0.001), 3 Hz (*r*=1.00, p<0.001; ICC_C-1_=1.00, p<0.001; ICC_A-1_=1.00, p<0.001), and 4 Hz (*r*=1.00, p<0.001; ICC_C-1_=1.00, p<0.001; ICC_A-1_=1.00, p<0.001).

### Toe tapping

Repeated measures ANOVA revealed a significant main effect of target frequency (F(3,27)=794.24, p<0.001), but no significant main effect of method (F(1,9)=4.86, p=0.06) or target frequency x method interaction (F(3,27)=2.18, p=0.11) on frequency of toe tapping (Figure 5A). Post-hoc analyses revealed significant differences in movement frequency between each pair of target frequencies (all p<0.001).

**Figure 5.**
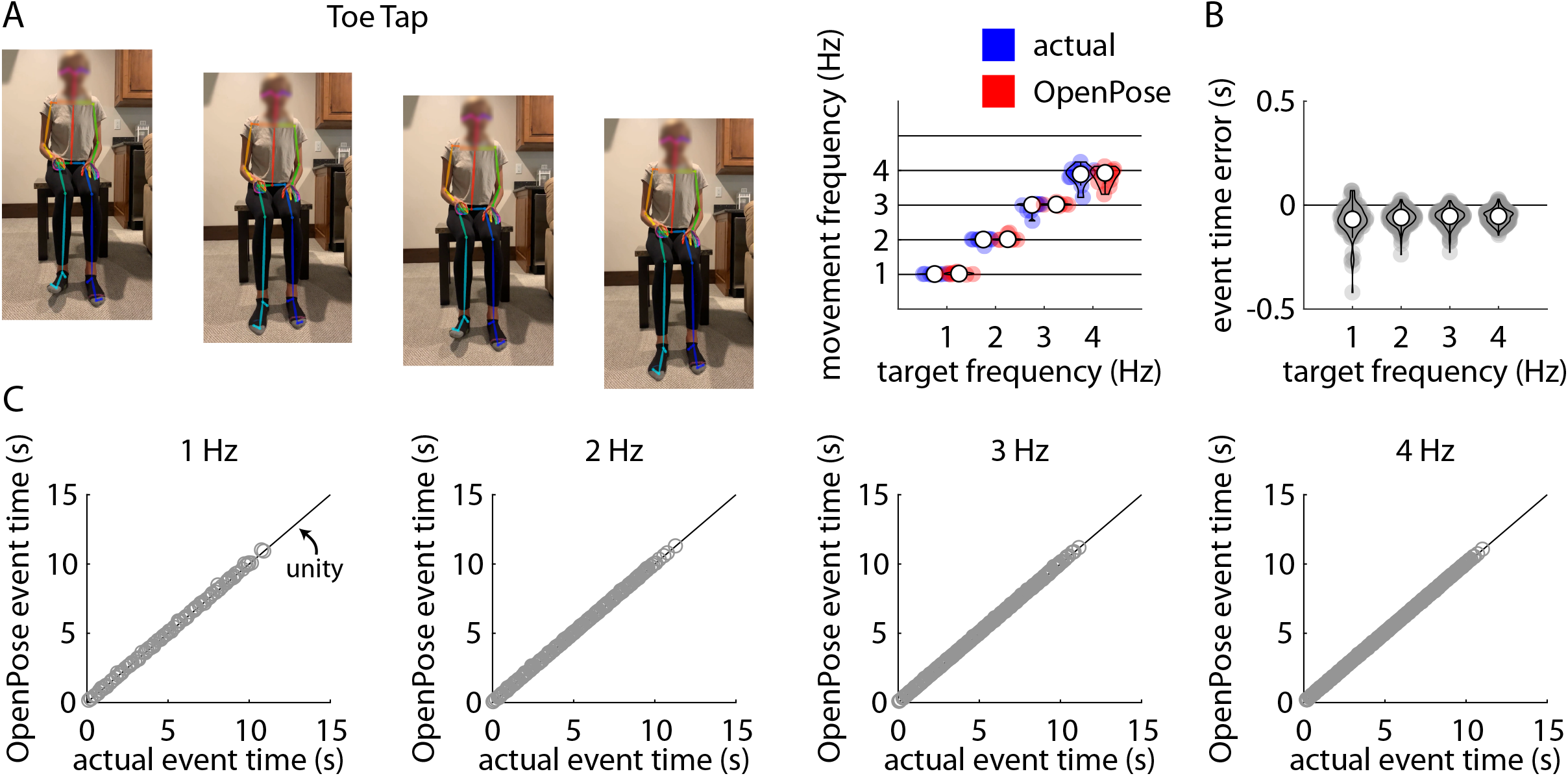
A) Representative example of stillframes from an OpenPose output video of a participant performing the toe tapping task (left) and violin plots showing comparisons of individual participant mean toe tapping frequencies measured manually (“actual”, blue) and using our pose estimation workflow (“OpenPose”, red) across all four target frequencies (right). B) Violin plots showing event time errors (i.e., differences between video timestamps of toe taps as measured manually and using pose estimation) for each individual toe tap across all participants. C) Scatter plots showing relationships between the video timestamps of all individual toe taps across all participants as measured manually (x-axis) or using our pose estimation workflow (y-axis) across all four target frequencies.

Across all trials, our pose estimation approach identified 989 toe taps and manual identification detected 1003 toe taps across all participants and trials; of these, there were 969 common toe taps detected by both methods. Distributions of errors between the pose estimation and manual detections of each toe tap detected by both methods are shown in Figure 5B. Figure 5C shows associations between manually-detected toe tap times and pose estimation-detected toe tap times for target frequencies of 1 Hz (*r*=1.00, p<0.001; ICC_C-1_=1.00, p<0.001; ICC_A-1_=1.00, p<0.001), 2 Hz (*r*=1.00, p<0.001; ICC_C-1_=1.00, p<0.001; ICC_A-1_=1.00, p<0.001), 3 Hz (*r*=1.00, p<0.001; ICC_C-1_=1.00, p<0.001; ICC_A-1_=1.00, p<0.001), and 4 Hz (*r*=1.00, p<0.001; ICC_C-1_=1.00, p<0.001; ICC_A-1_=1.00, p<0.001).

### Leg agility

Repeated measures ANOVA revealed a significant main effect of target frequency (F(3,27)=2005.84, p<0.001), but no significant main effect of method (F(1,9)=2.66, p=0.14) or target frequency x method interaction (F(3,27)=1.69, p=0.19) on frequency of heel tapping (Figure 6A). Post-hoc analyses revealed significant differences in movement frequency between each pair of target frequencies (all p<0.001).

**Figure 6.**
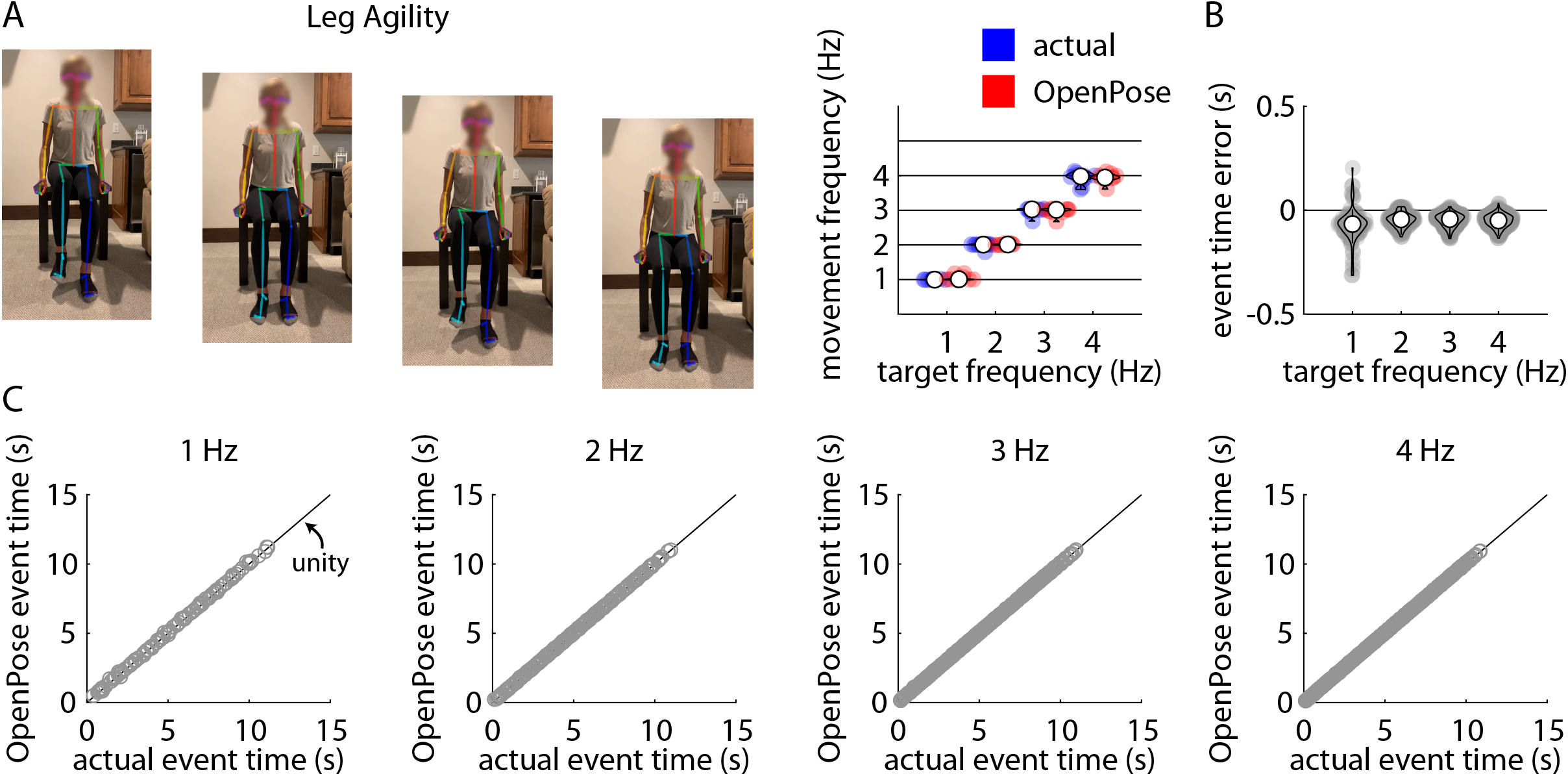
A) Representative example of stillframes from an OpenPose output video of a participant performing the leg agility (i.e., heel tapping) task (left) and violin plots showing comparisons of individual participant mean heel tapping frequencies measured manually (“actual”, blue) and using our pose estimation workflow (“OpenPose”, red) across all four target frequencies (right). B) Violin plots showing event time errors (i.e., differences between video timestamps of heel taps as measured manually and using pose estimation) for each individual heel tap across all participants. C) Scatter plots showing relationships between the video timestamps of all individual heel taps across all participants as measured manually (x-axis) or using our pose estimation workflow (y-axis) across all four target frequencies.

Across all trials, our pose estimation approach identified 1009 heel taps and manual identification detected 1001 heel taps across all participants and trials; of these, there were 996 common heel taps detected by both methods. Distributions of errors between the pose estimation and manual detections of each heel tap detected by both methods are shown in Figure 6B. Figure 6C shows associations between manually-detected heel tap times and pose estimation-detected heel tap times for target frequencies of 1 Hz (*r*=1.00, p<0.001; ICC_C-1_=1.00, p<0.001; ICC_A-1_=1.00, p<0.001), 2 Hz (*r*=1.00, p<0.001; ICC_C-1_=1.00, p<0.001; ICC_A-1_=1.00, p<0.001), 3 Hz (*r*=1.00, p<0.001; ICC_C-1_=1.00, p<0.001; ICC_A-1_=1.00, p<0.001), and 4 Hz (*r*=1.00, p<0.001; ICC_C-1_=1.00, p<0.001; ICC_A-1_=1.00, p<0.001).

## DISCUSSION

In this study, we demonstrated that pose estimation can be used to measure movement frequency across upper and lower extremity tasks in healthy adults. We used the pretrained human network included in the OpenPose demo to track the movements of healthy adults as they performed five different repetitive movement tasks (finger tapping, hand opening and closing, hand pronation and supination, toe tapping, and leg agility) at target frequencies spanning 1-4 Hz. We observed that our analysis workflow accurately detected movement events (e.g., finger taps, hand closures) and estimated movement frequencies that closely agreed with ground-truth measurements derived from manual inspection across all tasks and frequencies. Our results demonstrate that pose estimation can be used as a fast, inexpensive, video-based approach for quantitatively measuring human movement frequencies across upper and lower extremity tasks.

Interest in pose estimation has increased rapidly in the machine learning and neuroscience communities (9–14,27); however, clinical applications in humans are limited (18,22–24,28). The pros and cons of using currently available pose estimation algorithms for measurement of human movement have been discussed at length (29), and we will discuss our own impressions and suggestions below. Despite current limitations, this study and our prior work (21) have shown promising results when comparing pose estimation outputs to ground-truth measurements of clinically relevant movement parameters. We anticipate that the accuracy of human pose estimation algorithms will continue to improve, and we offer that this emerging technology has significant potential for versatile, quantitative, remote assessment of human motor function.

The repetitive movement tasks included in this study were selected because they are used in the MDS-UPDRS (3) to assess bradykinesia of the upper and lower extremities in persons with PD. This study is intended to be a preliminary step toward development of a quantitative, comprehensive, video-based approach for assessment of bradykinesia of the upper and lower extremities in persons with PD, and future work will be needed to assess the accuracy of movement frequency estimation in patient populations. Moreover, applications for PD in particular will need to expand upon our workflow to incorporate assessments of movement amplitude in addition to frequency.

To calculate movement frequencies, we detected event times manually (ground-truth) and using OpenPose keypoint trajectories. Across tasks, we tended to observe event time errors (i.e., differences between manual and pose estimation detections) of 1-2 video frames on average.

This suggests a small systematic bias in event detection between the two methods. However, we note that movement frequency can be estimated accurately since a phase shift (bias in event times) will preserve the true frequency despite differences in *when* the event times are detected. This is illustrated from our results demonstrating that OpenPose estimated movement frequencies well across a repertoire of motor tasks.

Other novel techniques that rely on wearables and mobile devices have been proposed to quantify performance of repetitive movement (4–8). While we show here that movement frequencies of tasks included in Part III of the MDS-UPDRS can be accurately obtained using video-based pose estimation in healthy adults, we did not test this approach against other device-based quantification methods. Indeed, other measurement devices may offer advantages that are difficult to quantify with our method. For example, wearable inertial measurement units (IMUs) may be more appropriate for free-living monitoring over longer durations. It is possible that different approaches could be integrated into a more comprehensive approach using different types of sensors to track various movement characteristics in PD. For quantitative information on repetitive movement in PD for clinical use, we propose that pose estimation may be favorable due to its ease of use, computational simplicity, and demonstrated accuracy.

We also found that our approach was quite versatile, as successful analysis of videos was achieved by variable methods of self-recording by participants, in variable environments, and with variable recording devices. Some participants placed a phone on a stable surface at the desired distance and height (as described in methods) and self-recorded without the assistance of another person. Others asked a family member or friend to record them according to these same guidelines, and both methods produced successful analyses. Additionally, participants recorded the videos in various locations and used their personal smartphones (the model varied among participants), and successful analyses were achieved for all participants. This underscores a significant advantage that this technique could have in clinical practice: all persons with a smartphone or tablet can likely record videos that could be successfully analyzed.

We did encounter a few difficulties using our approach and offer some potential solutions below. First, we chose to film the entire body and use both the OpenPose BODY_25 and Hand keypoint models because, in preliminary trials, this produced better hand keypoint detection than simply filming the hand and using the Hand keypoint model alone. Second, filming hand pronation/supination from the frontal plane (i.e., participant directly facing the camera with arms extended directly toward the camera) caused difficulty with consistent hand keypoint detection during pilot data collection. To address this issue, we asked participants to film all hand pronation/supination tasks in the sagittal plane (“from the side”). We found that this significantly improved hand keypoint tracking. A third observation was that, during finger tapping and hand opening and closing tasks, we achieved better tracking when the participant held the hand to the side of the body rather than extended in front of the torso (due to less overlap of the body keypoint tags with the hand keypoints). Finally, we observed that some postures (e.g., sitting with legs crossed) occasionally caused difficulty with BODY_25 keypoint tracking; therefore, we instructed participants to sit with “knees pointing forward and feet flat on the floor”.

## CONCLUSIONS

In this study, we used a pose estimation approach for fast, quantitative measurement of movement frequency during several upper and lower extremity tasks that are commonly included on neurological examinations. We found that this approach provided largely accurate identifications of event times and estimations of movement frequencies when compared to ground-truth measurements. Future work will aim to establish similar validity in clinical populations.

## ACKNOWLEDGMENTS

We acknowledge funding from the Association of Academic Physiatrists Rehabilitation Research Experience for Medical Students Fellowship to HLC and NIH grant R21AG059184 to RTR.

